# Correcting Cardiorespiratory Noise in Resting-state Functional MRI Data Acquired in Critically Ill Patients

**DOI:** 10.1101/2021.11.08.467753

**Authors:** Suk-tak Chan, William R. Sanders, David Fischer, John E. Kirsch, Vitaly Napadow, Yelena G. Bodien, Brian L. Edlow

**Author notes:** co-senior authors (Yelena G. Bodien, Brian L. Edlow). **Correspondence:** Suk-tak Chan, Address: Athinoula A. Martinos Center for Biomedical Imaging, Building 149, Thirteenth Street, Room 149-2301, Charlestown, MA 02129, United States.

## Abstract

Resting-state functional MRI (rs-fMRI) is being used to develop diagnostic, prognostic, and therapeutic biomarkers for critically ill patients with severe brain injuries. In studies of healthy volunteers and non-critically ill patients, prospective cardiorespiratory data are routinely collected to remove non-neuronal fluctuations in the rs-fMRI signal during analysis. However, the feasibility and utility of collecting cardiorespiratory data in critically ill patients on a clinical MRI scanner are unknown. We concurrently acquired rs-fMRI (TR=1250ms), cardiac and respiratory data in 23 critically ill patients with acute severe traumatic brain injury (TBI), and 12 healthy control subjects. We compared the functional connectivity results after denoising with cardiorespiratory data (i.e., RETROICOR) with the results obtained after standard bandpass filtering. Rs-fMRI data in 7 patients could not be analyzed due to imaging artifacts. In 6 of the remaining 16 patients (37.5%), cardiorespiratory data were either incomplete or corrupted. In both patients and control subjects, the functional connectivity corrected with RETROICOR did not significantly differ from that corrected with bandpass filtering of 0.008-0.125 Hz. Collectively, these findings suggest that there is a limited feasibility and utility to prospectively acquire high-quality cardiorespiratory data during rs-fMRI in critically ill patients with severe TBI for physiological correction.

## 1 INTRODUCTION

Over the past decade, resting-state functional magnetic resonance imaging (rs-fMRI) studies have demonstrated the potential diagnostic and prognostic utility of brain network mapping in patients with a broad range of brain injuries, including traumatic brain injury (TBI) (Guo et al., 2019; Kondziella et al., 2017; Peran et al., 2020; Threlkeld et al., 2018), hypoxic-ischemic injury (HIE) from cardiac arrest (Koenig et al., 2014; Norton et al., 2012; Peran et al., 2020; Pugin et al., 2020; Sair et al., 2018), and hypoxic injury from coronavirus disease 2019 (COVID-19) (Fischer et al., 2020). In addition, rs-fMRI is now used as a pharmacodynamic biomarker in early-stage clinical trials to assess brain responses to targeted therapies aimed at promoting recovery of consciousness (Edlow et al., 2020). Nevertheless, despite the growing evidence that rs-fMRI has potential as a diagnostic, prognostic, and therapeutic biomarker, the optimal data acquisition, pre-processing, and analysis methods for rs-fMRI have not been determined for critically ill patients with acute brain injuries. For example, a fundamental pre-processing question is how to account for non-neuronal, physiological fluctuations, such as cardiac pulsation and respiration. These cardiorespiratory signal fluctuations contribute noise to the data, obscuring the blood-oxygen-level dependent (BOLD) signals used to identify functional network connectivity. Accounting for these cardiorespiratory signal fluctuations during pre-processing is particularly relevant in critically ill patients, who may experience fluctuations in vital signs during the rs-fMRI scan, causing potentially unpredictable changes in the BOLD signal. As a result, the underlying neuronal dynamics measured by functional network connectivity may be masked.

Prior studies in healthy human subjects have shown that, compared to standard bandpass filtering, prospective acquisition of cardiorespiratory data may improve the signal-to-noise properties of spontaneous BOLD fluctuations (Chen et al., 2020; Chu et al., 2018; Glover et al., 2000; Murphy et al., 2013), because these cardiorespiratory data facilitate signal correction based on subjects’ unique cardiac and respiratory oscillations. In addition, cardiac pulsation and respiration signals have different frequencies and amplitude characteristics (Gray et al., 2009), and therefore may not be amenable to correction with a single bandpass filter. Notably, most studies that prospectively collected cardiorespiratory data enrolled healthy control subjects who had relatively stable hemodynamics, are able to tolerate long scan times, and were scanned in research settings with precise instrumentation (Chan et al., 2020; J. Lee et al., 2019). Conversely, setting up additional equipment and optimizing the cardiorespiratory signals prolong scan time, posing safety risks for critically ill patients. Patients in the intensive care unit (ICU) must be scanned on a clinical scanner using standard physiological monitoring instrumentation integrated with the MRI system, where the cardiorespiratory data acquired may be suboptimal for rs-fMRI signal denoising. Moreover, the quality of cardiorespiratory data also depends on a patient’s injury burden, cardiopulmonary stability, hemodynamic fluctuations, and cooperation. Involuntary motion is common in critically ill patients with altered consciousness (Bodien et al., 2017; Edlow et al., 2017; Threlkeld et al., 2018), and the placement of physiological sensors can be complicated by concurrent injuries to the chest, abdomen, and extremities. Indeed, these safety, scientific and clinical challenges likely explain why few fMRI studies have been performed on critically ill patients (Bodien et al., 2017), and why there have been no studies, to our knowledge, acquiring cardiorespiratory data specifically for the pre-processing of rs-fMRI data in critically ill paitents.

In this prospective, observational study, we investigated the feasibility of acquiring cardiorespiratory data during rs-fMRI in critically ill patients with acute severe TBI and the utility of using these data to correct for non-neuronal signals in the BOLD data. The study was performed on a 3 Tesla MRI scanner located in the Neurosciences ICU (NeuroICU) at an academic medical center. We hypothesized that cardiorespiratory data acquisition is feasible in a clinical setting for critically ill patients and that it reduces physiological confounds more effectively than does bandpass filtering. In the feasibility analysis, we assessed the success rate of acquiring technically useful, uncorrupted cardiac and respiratory data. In the utility analysis, we compared the functional connectivity brain maps corrected with RETROICOR (Glover et al., 2000) to those corrected with standard bandpass filters. Recognizing that there is no gold-standard for “ground-truth” physiological correction of functional brain connectivity, we tested the hypothesis that RETROICOR correction using cardiorespiratory signals yields higher levels of network connectivity than does standard bandpass filtering.

## 2 MATERIALS AND METHODS

### 2.1 Participants

We enrolled 12 healthy volunteers and 23 patients with acute severe TBI, defined by a post-resuscitation Glasgow Coma Scale (GCS) score ≤ 8 prior to admission to an ICU. All the patients were enrolled consecutively from the NeuroICU, surgical ICU, or multi-disciplinary ICU at Massachusetts General Hospital (MGH), as part of an ongoing observational study (ClinicalTrials.gov NCT03504709). Healthy control subjects had no history of brain injuries, neurological disease, psychiatric disease, cardiovascular disease, or any history of diabetes, hypertension, or renal disease, and were recruited by e-mail and poster placement within the MGH hospital network.

All components of this study were performed in compliance with the Declaration of Helsinki, and all procedures were approved by the hospital’s Human Research Committee. Written informed consent was obtained from healthy subjects and patients’ surrogate decision-makers.

### 2.2 MRI Acquisition

We performed the MRI on a 3 Tesla Skyra scanner (Siemens Medical, Erlangen, Germany) in the NeuroICU with a 32-channel head coil. Foam pads and inflatable positioning pads were used to minimize head motion. We acquired the following MRI datasets on each subject: 1) high-resolution sagittal images acquired with volumetric T1-weighted 3D-MEMPRAGE sequence (TR=2530ms, TE=1.69ms/3.55ms/5.41ms/7.27ms, flip angle=7°, FOV=256×256mm, matrix=256×256, slice thickness=1mm); 2) BOLD-fMRI images acquired with echo-planar imaging (EPI) sequence (TR=1250ms, TE=30ms, flip angle=65°, FOV=212×212mm, matrix=106×106, slice thickness=2mm, slice gap=0mm, duration=10 minutes) while the subject was at rest. Subjects were instructed to keep their eyes open during the rs-fMRI scans. We aimed to acquire two rs-fMRI datasets for each subject – one at the beginning of the MRI scan, and one at the end of the scan – separated by approximately 30 minutes of stimulus-based and task-based fMRI paradigms.

We optimized the MRI sequence with a short TR of 1.25 seconds and a resting-state scan length of 10 minutes. This made the maximum sampling frequency 0.4 Hz, allowing the removal of most of the fluctuations due to respiratory cycles. The narrow frequency spacing of about 0.002 Hz based on the scan duration offered proper sampling of signal changes down to 0.002 Hz and more precise removal of fluctuations when bandpass filtering strategy was applied.

We recorded the time series of both optical plethysmography and respiration using the Siemens Physiological Monitoring Unit (Siemens Healthcare, Erlangen, Germany). Optical plethysmography was measured instead of electrocardiogram for cardiac pulsation because the optical signals were less contaminated by the imaging gradient changes. Breath-by-breath respiratory cycles were measured by pressure changes in the pneumatic bladder located between the skin surface and the respiratory belt around the upper abdomen. All cardiorespiratory data recordings were synchronized using the timestamps in the data recordings and image headers.

### 2.3 Data Analysis

The imaging and cardiorespiratory data from the first rs-fMRI scan for each subject were used in the following analyses, except for one patient (P3) because the first rs-fMRI scan was terminated in the middle due to a technical issue with the head coil resulting in signal void in the anterior part of the brain. For this patient, the imaging and cardiorespiratory data from the second rs-fMRI scan were used.

#### 2.3.1 Processing of Cardiorespiratory Data

We used Matlab R2020a (Mathworks, Inc., Natick, MA, USA) to analyze the cardiorespiratory data. The peaks on the optical plethysmography time series serve as a surrogate of R peaks in electrocardiogram, while the peaks and troughs on the respiratory time series indicated end inspiration and end-expiration respectively (**Supplementary Figure 1**). Peaks and troughs on the time series of optical plethysmography and respiration were determined using a custom Matlab function (Chan et al., 2020) and corrected on the graphical user interface. The cardiac phase used in RETROICOR (Glover et al., 2000) advances linearly from 0 to 2π during each R-R interval and is reset to 0 for the next cycle. The inspiratory phase spans from 0 to π and the expiratory phase spans from 0 to −π.

#### 2.3.2 Pre-processing of Resting-State BOLD-fMRI Data

All BOLD-fMRI data were imported into the Analysis of Functional NeuroImage (AFNI) software (Cox, 1996) (National Institute of Mental Health, http://afni.nimh.nih.gov) for pre-processing. The first 12 volumes of each functional dataset, collected before equilibrium magnetization was reached, were discarded. Artifactual spikes were removed from the time series in each voxel using ‘3dDespike’ in AFNI. In order to compare resting-state connectivity after RETROICOR versus standard bandpass filtering, the resting-state BOLD data were separately processed with two processing pipelines: RETROICOR (RETROICOR Pipeline) and standard bandpass filtering (BANDPASS Pipeline). We focused on RETROICOR, rather than other physiological correction methods that employ a component-based approach (e.g., CompCor), because some of these methods require accurate specification of a “noise” region-of-interest in white matter or cerebrospinal fluid (Behzadi et al., 2007). In patients with distorted brain anatomy and BOLD signal changes in white matter due to traumatic microbleeds (S. Lee et al., 2018), defining a precise “noise” region-of-interest is especially challenging. A schematic diagram of our data analysis algorithm is shown in **Figure 1**.

**Figure 1.**
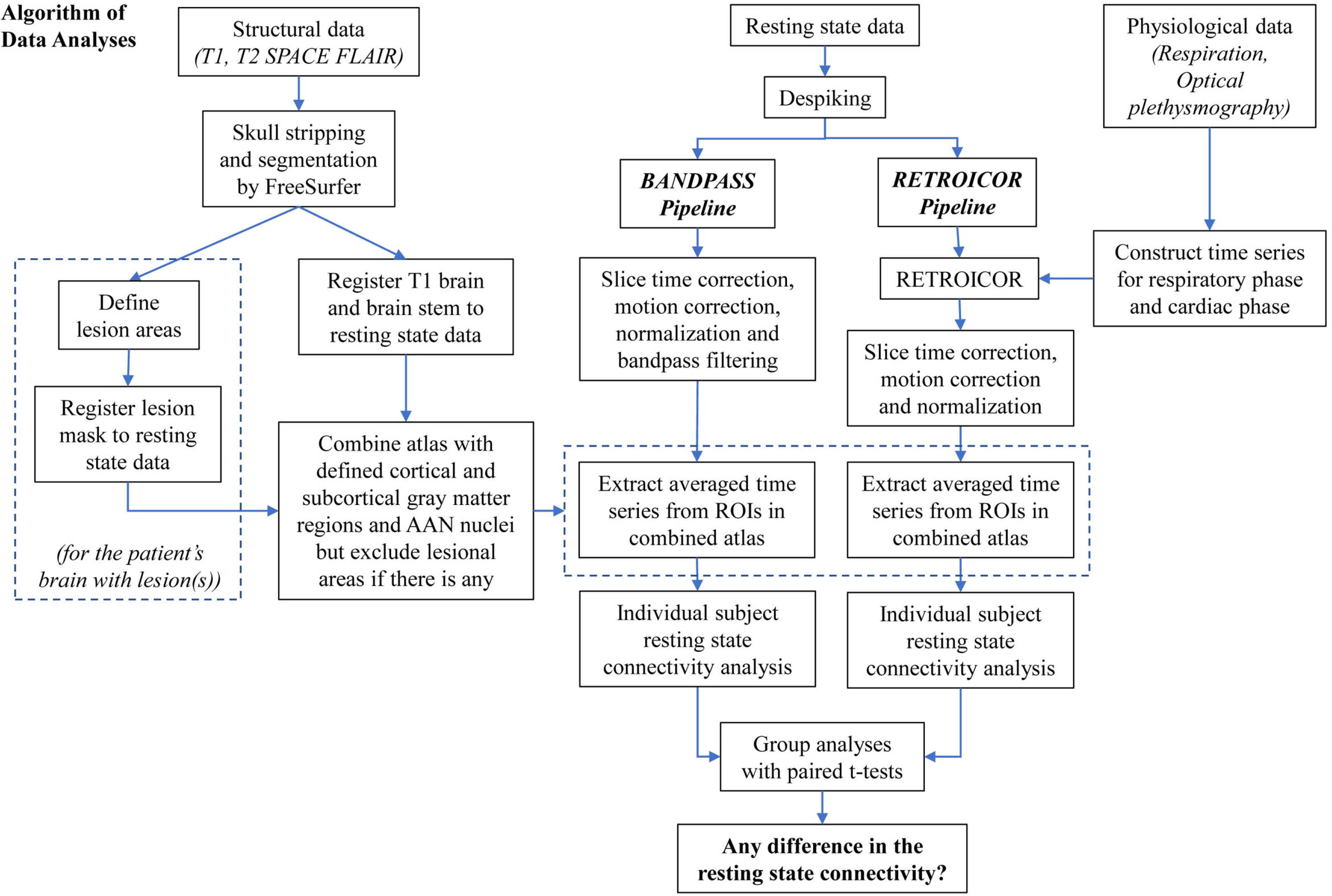
Algorithm of data analyses.

##### 2.3.2.1 RETROICOR Pipeline

Time series of cardiac and respiratory phases were imported into the program ‘3dretroicor’ in AFNI to remove cardiac and respiratory motions of the resting state BOLD-fMRI after despiking. Each functional dataset in the RETROICOR Pipeline was corrected for slice timing, motion-corrected, and co-registered to the first image of the functional dataset using three-dimensional volume registration. Voxels located within the ventricles and outside the brain were defined by the parcellated brain volume using FreeSurfer (Dale et al., 1999; Fischl et al., 1999) (MGH/MIT/HMS Athinoula A. Martinos Center for Biomedical Imaging, Boston, http://surfer.nmr.mgh.harvard.edu) and were excluded from subsequent analyses. In the co-registered functional dataset, motion outliers were defined as any timepoint with 5% of brain voxels having the averaged derivative change of translational and rotational motion parameters of more than 0.4. The coregistered dataset was then normalized to its mean intensity value across the time series for the percent BOLD signal changes (ΔBOLD). In the normalized functional dataset, the time series of each voxel was detrended with the fifth order of polynomials to remove the low drift frequency (<0.005 Hz). The low drift frequency and the components of motion were removed in one single process using orthogonal projection. Individual subject brain volumes with time series of ΔBOLD were used in the connectivity analysis.

##### 2.3.2.2 BANDPASS Pipeline

Slice timing correction, motion correction, co-registration, normalization, and removal of low drift frequency and motion components were applied to each functional dataset after despiking in the BANDPASS pipeline using the same approach as in the RETROICOR pipeline. Bandpass filters of different frequency bandwidths were separately applied to the clean functional dataset. The starting frequency of the bandpass filters was 2^x^, where x ranged from −7 (i.e., 2^−7^=0.008 Hz) to −2 (i.e., 2^−2^=0.25 Hz) at an increment of the exponential of +1. The ending frequency of the bandpass filters was 2^y^, where y ranged from −6 (i.e., 2^−6^=0.016 Hz) to −1 (i.e., 2^−1^=0.5 Hz) at an increment of the exponential of +1. A total of twenty-one frequency bandwidths were used as bandpass filters and 21 filtered datasets were obtained.

#### 2.3.3 Individual Subject Resting-State Connectivity Analysis

For each subject, we constructed a brain mask with 82 cortical and subcortical gray matter brain regions parcellated by the software FreeSurfer (Dale et al., 1999; Fischl et al., 1999) and 16 brainstem nuclei from the Harvard Ascending Arousal Network (AAN) atlas (Edlow et al., 2012). No single voxel was included in more than one brain region. If a subject had a large brain lesion, such as a hemorrhagic contusion, visible on the T1-weighted structural images, the study investigator (STC) constructed a lesion mask by manually drawing the lesion on the high-resolution T1-weighted images as a region-of-interest using FreeSurfer (http://surfer.nmr.mgh.harvard.edu), as done previously (Diamond et al., 2020). The brain atlas with defined cortical and subcortical gray matter regions and AAN nuclei and the brain lesion masks were registered to brain volumes with time series of ΔBOLD via the first brain volume in the motion-corrected functional dataset. Brain regions that overlapped with brain lesion masks were excluded from the connectivity analysis.

For the normalized dataset in the RETROICOR Pipeline, the averaged time series of ΔBOLD in each brain region were correlated with that of each other brain region using the program ‘3dNetCorr’ (Taylor & Saad, 2013). Pearson’s correlation coefficients were calculated from 3403 region pairs. In the BANDPASS Pipeline, the same correlation analysis was applied to each normalized dataset. For both pipelines, the Pearson’s correlation coefficient from each region pair was transformed to a Fisher’s z value, to indicate the connectivity strength for the group analysis.

#### 2.3.4 Group Analysis

Age as a continuous variable was summarized as median and interquartile range, and group (TBI and control) difference was assessed using the Kruskal-Wallis test. Sex as a categorical variable was summarized as frequencies; group difference was assessed using the Chi-square test.

The connectivity strength indicated by Fisher’s z values from patient and control groups were analyzed separately. The potential difference in the brain connectivity for each region pair was first explored using paired t-test to compare the Fisher’s z values derived from the RETROICOR Pipeline and those from the commonly used bandpass filter of 0.008-0.125 Hz in the BANDPASS Pipeline. Such a comparison was repeated for 3403 region pairs. False discovery rate was used to correct the multiple comparisons. A significant difference was considered at p_fdr_<0.05.

To further explore if the frequency bandwidth of 0.008-0.125 Hz was optimal for bandpass filtering of rs-fMRI data, we used Pearson’s correlation to measure the correlation of averaged connectivity strength between the RETROICOR Pipeline and 21 bandpass filters in the BANDPASS Pipelines. For each region pair, the mean Fisher’s z value was calculated separately from the RETROICOR Pipeline and each bandpass filter in the BANDPASS Pipeline in the same group of subjects. The mean Fisher’s z values from 3403 region pairs in two processing arms were correlated. A significant correlation was considered at p<0.05.

Intraclass correlation was used to study the similarity of the connectivity strength between the two pipelines in each region pair. The intraclass correlation coefficients between the two pipelines for each region pair were calculated between the Fisher’s z values from the RETROICOR Pipeline and each bandpass filter in the BANDPASS Pipeline in the same group of subjects. False discovery rate was used to correct for multiple comparisons. A significant correlation was considered at p_fdr_<0.05.

## 3 RESULTS

A total of 35 subjects were enrolled (age range 18 to 78 years). Twenty-three were patients (median=37.0 years, interquartile range IQR=27.5-63.5years; 15M), and 12 were healthy controls (median=32.5 years, interquartile range IQR=28.8-35.8 years; 3M). The schematic diagram of subject inclusion and exclusion is shown in **Supplementary Figure 2**. The head impact mechanism and the level of consciousness at the time of the rs-fMRI scan for patients are shown in **Table 1**. The distribution of brain lesions in the patients is shown in **Supplementary Figure 3**. Lesions in most patients occurred in inferior frontal and anterior temporal areas. Of the 23 enrolled patients, 7 patients were excluded from subsequent data analyses due to failed brain segmentation caused by brain distortion, large MR signal void (e.g., from ventriculoperitoneal shunts), or gadolinium contrast that was administered for clinical imaging before the research MRI sequences (see **Table 1 and Supplementary Figure 2**). No healthy subjects were excluded due to image quality.

**Table 1.**
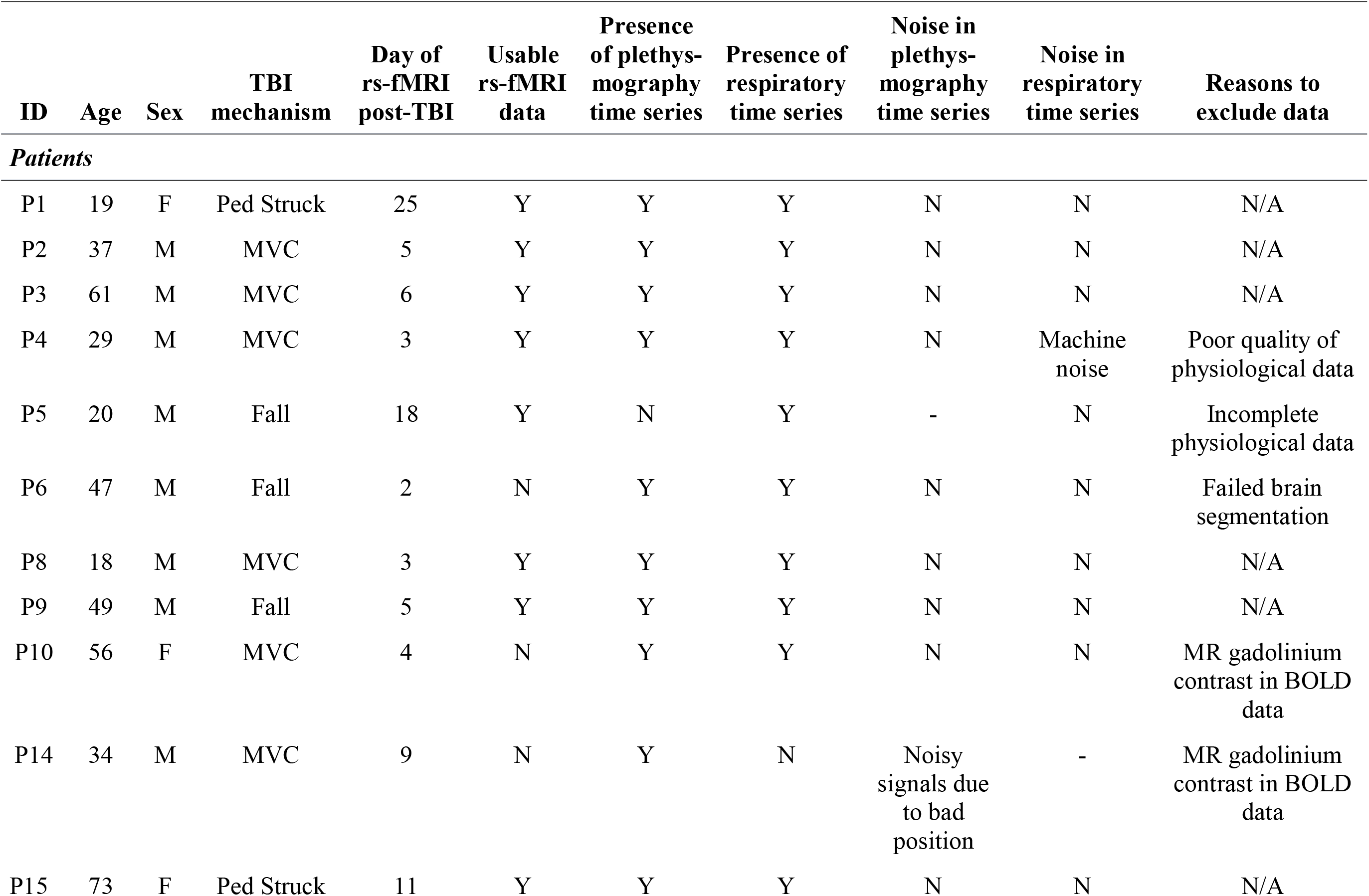
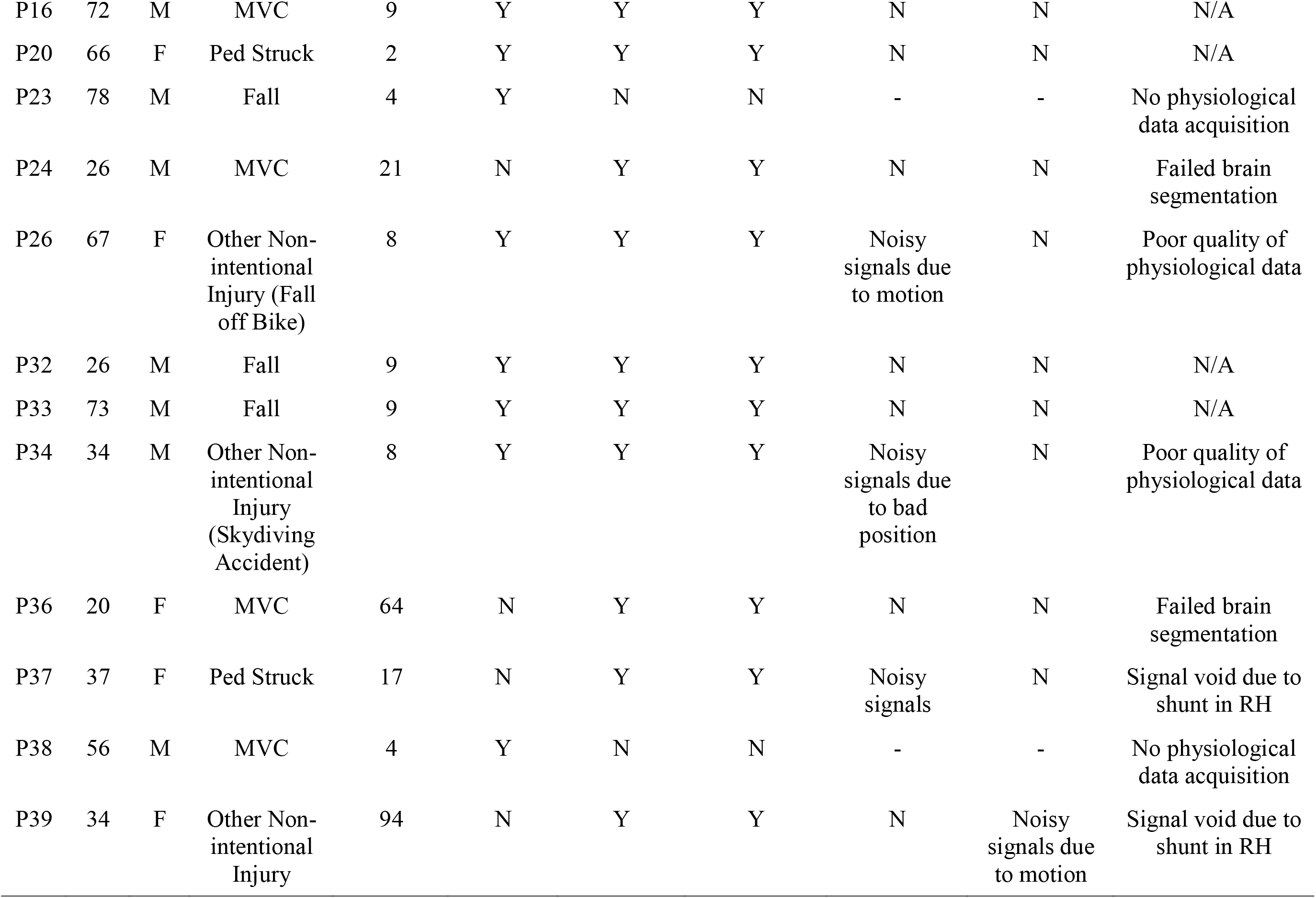
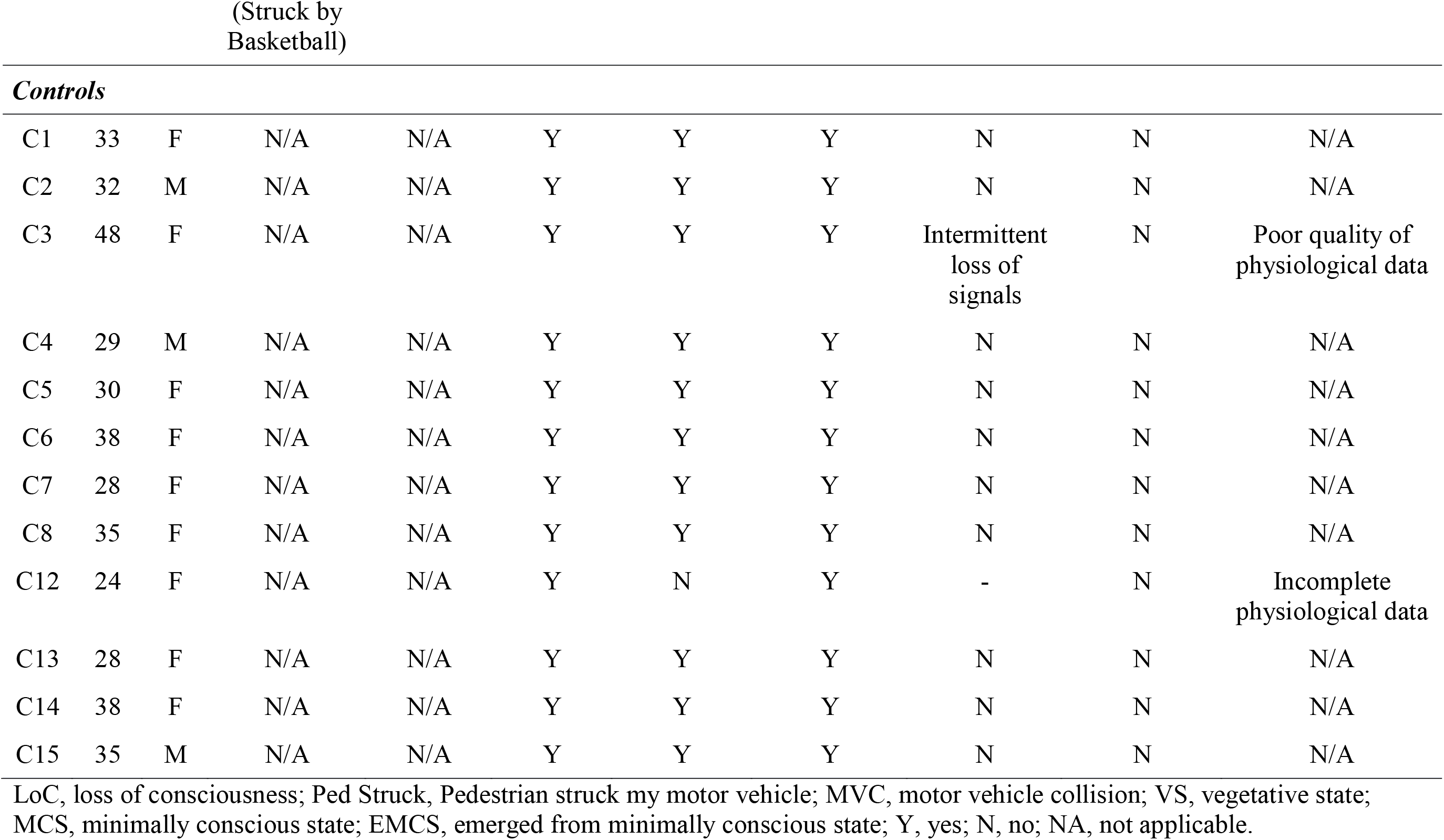
Feasibility of physiological monitoring during resting-state functional MRI in critically ill patients and controls.

**Table 2.** Summary of subject demographics.

|  | **Controls** | **Patients** | ***p*** |
| --- | --- | --- | --- |
| N | 10 | 10 |  |
| Age | 32.5 [29.2,35.0] | 55.0 [28.8,70.5] | 0.173 |
| Sex: Male | 3 (30.0) | 7 (70.0) | 0.180 |

Categorical variable (i.e., sex) is summarized as frequency (percentage) and continuous variable (i.e., age) is summarized as median [interquartile range]. No significant difference in age and sex was found between patients and controls.

### 3.1 Feasibility of Acquiring Data for Prospective Physiological Correction

**Table 1** provides details about why subject data were excluded from the group analyses, and a summary is shown in **Supplementary Figure 2**. Of 28 subjects with usable fMRI data (16 patients, 12 healthy controls), neither cardiac nor respiratory data could be acquired in 2 patients because extra time had been used to prepare these two patients for MRI scan, and the MRI scan had to be completed within a limited period of time. The time constraint in data acquisition also resulted in incomplete cardiorespiratory data acquisition (i.e., missing either cardiac or respiratory data) in 1 patient and 1 healthy control. The quality of plethysmography data was poor in 3 patients and 1 healthy subject due to subject motion, suboptimal sensor position and the intermittent loss of signals that magnetic interferences may cause on physiological signal reception in the scanner.

Examples of poor quality plethysmography data are shown in **Supplementary Figure 4**. In total, 14% (4/28) of all subjects had incomplete cardiorespiratory data, and another 14% (4/28) of subjects had poor quality cardiorespiratory data. Thus, 63% of patients with intact fMRI data (10/16) and 83% of healthy controls (10/12), had complete and usable cardiac and respiratory data for physiological correction.

Datasets from those 10 patients (median=55.0 years, interquartile range IQR=33.1-65.7 years; 7M) and 10 healthy controls (median=32.5 years, IQR=29.9-35.3 years; 3M) were included in the subsequent utility analyses. No significant difference in age (p=0.17) or sex (p=0.18) was found between patients and controls.

### 3.2 Utility of Cardiac and Respiratory Data for Physiological Correction

#### 3.2.1 Comparison of connectivity strength after RETROICOR correction and bandpass filtering of 0.008-0.125 Hz

Brain connectivity of a representative patient and a control subject after RETROICOR correction and bandpass filtering of 0.008-0.125 Hz are shown in **Supplementary Figure 5**. There is no noticeable difference in brain connectivity between the two methods of physiological noise correction (RETROICOR versus BANDPASS Pipelines) in the patient or control subject.

No significant difference was found in the group comparison of connectivity strength after RETROICOR correction and bandpass filtering of 0.008-0.125 Hz in both patients and healthy subjects (p_fdr_>0.05) **(Figure 2)**.

**Figure 2.**
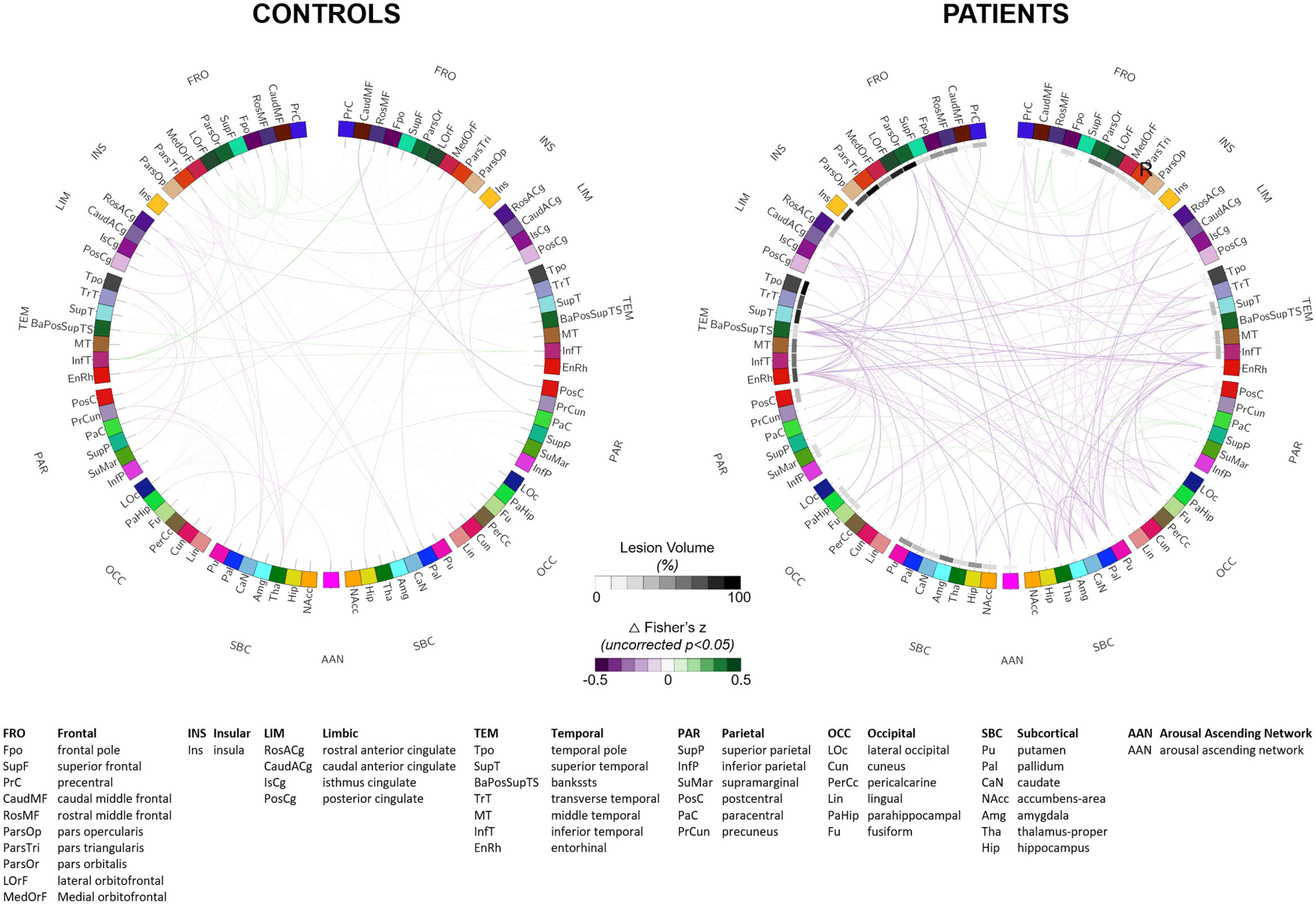
Comparison of brain connectivity between RETROICOR correction and BANDPASS of 0.008-0.125Hz in controls (n=10) and patients (n=10). No significant difference between RETROICOR and BANDPASS correction was found after correcting for multiple comparisons (p_fdr_>0.05).

#### 3.2.2 Comparison of connectivity strength after RETROICOR correction and bandpass filtering of 21 different frequency bandwidths

The mean Fisher’s z values derived from RETROICOR correction and bandpass filtering in different region pairs started to attain the highest correlation at the bandpass filtering of 0.008-0.125 Hz in patient group (Pearson’s r = 0.958, p<0.001) and control group (Pearson’s r = 0.981, p<0.001) (**Figures 3A and 3B**). The correlation stayed relatively high at the bandpass filtering of 0.008-0.25 Hz (Patients: Pearson’s r = 0.957, p<0.001; Controls: Pearson’s r = 0.981, p<0.001) and 0.008-0.5 Hz (Patients: Pearson’s r = 0.957, p<0.001; Controls: Pearson’s r = 0.981, p<0.001).

**Figure 3.**
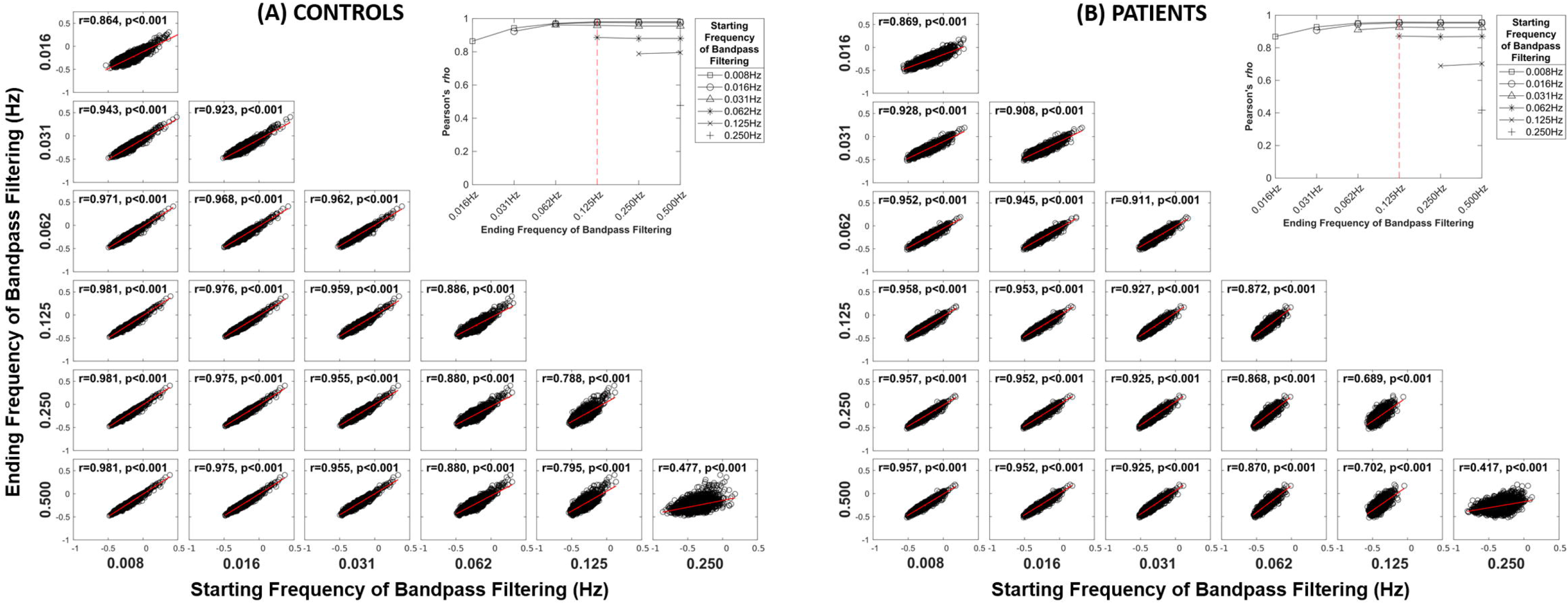
Correlation analyses of brain connectivity between RETROICOR and BANDPASS pipelines in controls (n=10) (A) and patients (n=10) (B). The Pearson’s r and the p values are shown in each correlation analysis. The top three frequency bandwidths in BANDPASS pipeline showing the highest correlation with RETROICOR pipeline are bound in red boxes. The curves at the right upper corner show the changes of correlation coefficients in the comparisons between the two pipelines. Broken red line indicates the ending frequency of bandpass filtering in BANDPASS pipeline when the highest correlation attains.

An increased number of region pairs had a significant intraclass correlation coefficient between RETROICOR and bandpass filtering with the starting frequency at 0.008 Hz in both patients and control (**Figure 4**). The top three frequency bandwidths with the highest number of region pairs showing significant intraclass correlation are 0.008-0.125 Hz, 0.008-0.25 Hz, and 0.008-0.5 Hz.

**Figure 4.**
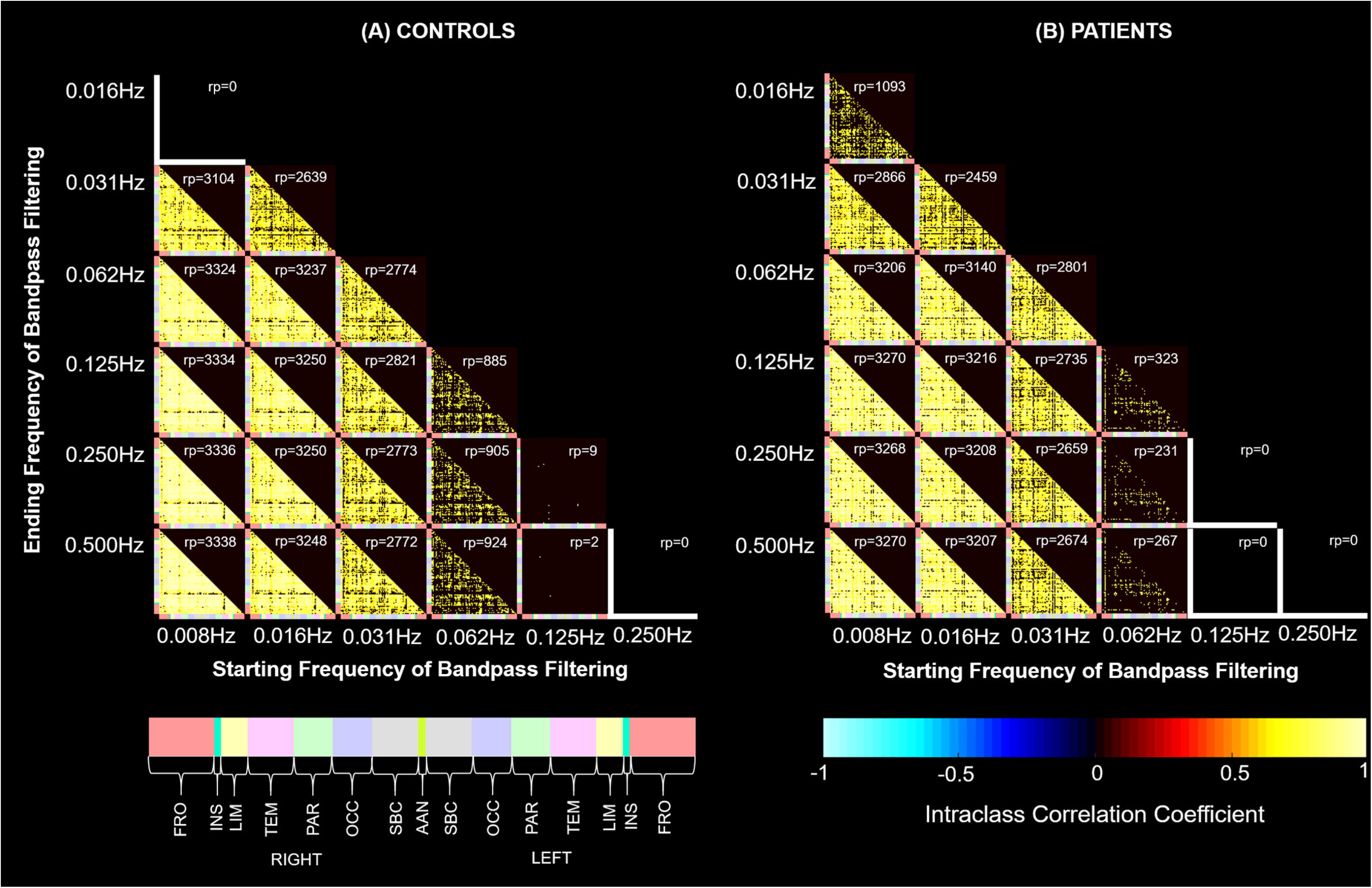
Intraclass correlation of connectivity strength between RETROICOR and BANDPASS pipelines in controls (n=10) (A) and patients (n=10) (B). The 83 brain regions on the x-axis and y-axis of each matrix are grouped into cerebral lobes indicated by colors. The number of region pairs (rp) showing significant intraclass correlation after correcting for multiple comparisons are shown at the upper right hand corner of each matrix. FRO, frontal; INS, insula; LIM, limbic; TEM, temporal; PAR, parietal; OCC, occipital; SBC, subcortical; ANN, arousal ascending network.

## 4 DISCUSSION

We investigated the feasibility and utility of cardiorespiratory noise correction in rs-fMRI data acquired in critically ill patients. In 6 of 16 patients, we could not conduct prospective cardiorespiratory noise correction due to missing or corrupted physiological data. Furthermore, we found no significant differences in functional brain connectivity for patients or controls when we compared two physiological noise correction methods (RETROICOR using prospectively collected cardiac and respiratory data versus bandpass filtering at 0.008-0.125 Hz). Collectively, these findings suggest that prospective cardiorespiratory data acquisition during rs-fMRI has limited feasibility and utility for correcting physiological noise in rs-fMRI data acquired in critically ill patients with acute severe TBI.

### 4.1 Impact of physiological correction using respiratory and cardiac data in ICU settings

Correction of cardiorespiratory noise that may confound rs-fMRI connectivity data is essential to ensure that BOLD signal modulation and the reduction in the degrees of freedom are identical across subjects (Murphy et al., 2013). A common approach to correct for cardiorespiratory noise is to regress prospectively collected cardiac and respiratory signals from the rs-fMRI data using RETROICOR. In our study, the advantage of using individualized cardiorespiratory noise correction via RETROICOR was outweighed by the clinical limitations of acquiring cardiac and respiratory data in critically ill patients. Excellent quality cardiac and respiratory data are required for physiological correction using RETROICOR. However, patients with severe acute TBI often have reduced tolerance for lying supine in the scanner, unstable hemodynamics, peripheral injuries, and impaired awareness. Futhermore, patients with critical illness must be scanned on a clinical MRI system that typically lacks precise instrumentation for acquiring cardiac and respiratory data. In our study, these safety considerations and logistical factors made it infeasible to consistently collect the high-quality cardiac and respiratory data required for RETROICOR.

In the absence of prospectively collected cardiac or respiratory data, band-pass filtering, typically at 0.008-0.125 Hz, may be used to correct for physiological noise. Although bandpass filtering is associated with a loss of degrees of freedom due to the removal of frequency bins (Murphy et al., 2013), this approach ensures that data can be analyzed in a standardized manner across all individuals. We found that brain connectivity metrics derived from RETROICOR and bandpass filtering were similar. We also tested multiple ranges of bandpass filtering and found that brain connectivity corrected with bandpass filtering of 0.008-0.125 Hz had the strongest correlation with brain connectivity corrected with RETROICOR. This bandpass filtering range has been used in many prior studies (Hillary et al., 2011; Huang et al., 2014; Threlkeld et al., 2018) even though it is associated with a loss of degrees of freedom, in our study from 200 to 60.

### 4.2 Correcting spontaneous fluctuations at 0.125 Hz or higher in critically ill patients

Our study focused on correcting physiological signals due to respiratory (Brosch et al., 2002; Raj et al., 2001) and cardiac cycles (Dagli et al., 1999) with frequencies of 0.15 Hz or higher. We did not correct for spontaneous fluctuations below 0.125 Hz because many physiological fluctuations below 0.125 Hz are related to neuronal signaling, especially in critically ill patients. Fluctuations related to intracranial pressure (0.008-0.03Hz) (Lundberg, 1960), respiratory gas exchange (0.008-0.03Hz) (Chan et al., 2020; Lenfant, 1967), respiratory variation (∼0.03Hz) (Birn et al., 2006; Chang et al., 2013), end-tidal carbon dioxide fluctuations (0-0.05Hz) (Wise et al., 2004), variation in arterial pressure (0.05-0.15Hz) (Mayer, 1876; Obrig et al., 2000), or heart rate variability (0.05-0.15Hz) (Chang et al., 2013) **(Supplementary Figure 6)** all occur at a similar frequency to spontaneous BOLD fluctuations. Among these fluctuations, respiratory gas exchange (Chan et al., 2020), respiratory variation (Birn et al., 2006), end-tidal carbon dioxide fluctuations (Wise et al., 2004), and heart rate variability (Chang et al., 2013) were previously found to be correlated with oscillations of the default mode network (DMN). Therefore, extra caution is required when removing the physiological “noise” in the frequency bandwidth below 0.125Hz to avoid removing important information about functional connectivity.

### 4.3 rs-fMRI in the intensive care unit setting

Resting-state fMRI has historically been used as an investigational tool to study functional brain network connectivity in patients with a broad range of neurological and psychiatry disorders. In critically ill patients with severe brain injuries, emerging evidence suggests that DMN connectivity is associated with recovery of consciousness (Threlkeld et al., 2018), and DMN and salience network connectivity predict long-term functional outcomes (Sair et al., 2018). Rs-fMRI is also being used as a pharmacodynamic biomarker in treatment studies aimed at promoting recovery of consciousness (Edlow et al., 2020). More recently, rs-fMRI has emerged as a promising clinical tool (Boerwinkle et al., 2020) for mapping functional brain connectivity and relating it to the capacity for recovery of awareness (Threlkeld et al., 2018). In the early stages of clinical rs-fMRI implementation, it is essential that data acquisition and processing are standardized, and that safety and feasibility are optimized. Our findings support this goal and inform future clinical implementation efforts by providing initial evidence that cardiorespiratory data are difficult to collect uniformly and have limited utility for evaluating functional connectivity.

### 4.4 Limitations of this study

Our study is limited a small sample size, which reflects the difficulty of transporting and scanning critically ill patients in the early days of recovery from severe TBI. A larger sample size may have revealed subtle differences between RETROICOR and bandpass methods. In addition, we used the plethysmography and chest recordings acquired with the standard physiological monitoring unit in the MRI scanner, which is designed for clinical monitoring and pacing image acquisition for cardiac and abdominal imaging. The physiological monitoring unit was not designed for the quantitative assessment of cardiac and respiratory activity. Although a larger sample and more precise equipment may have lead to different findings, our goal was to assess the feasibility and utility of correcting cardiorespiratory noise in an ICU setting that reflects the practical challenges of acquiring fMRI data in patients with acute severe TBI. In this context, our findings indicate that as rs-fMRI is integrated into the clinical assessment of patients with severe TBI, bandpass filtering at 0.008-0.125Hz can be used as an alternative to prospective acquisition of cardiorespiratory data for physiological correction in the functional connectivity analysis.

## 5 CONCLUSION

We found that prospective cardiorespiratory correction has limited feasibility and utility in critically ill patients. Given currently available technology and logistics of cardiorespiratory data acquisition, our findings suggest that studies using cardiorespiratory correction in critically ill patients may need to exclude a significant number of patients and accept a reduced sample size. We also observed that correction using prospective cardiorespiratory data acquisition may not provide a significant advantage of analytic utility over retrospective bandpass filtering correction in critically ill patients.

These observations suggest that physiological correction of rs-fMRI using prospective acquisition of cardiorespiratory data has limited feasibility and utility in rs-fMRI studies of critically ill patients.

## 6

**Supplementary Figure 1**. Example of cardiac and respiratory phases derived from cardiac and respiratory data respectively for RETROICOR (Glover et al., 2000). Orange lines indicate the cardiac and respiratory phases used in RETROICOR. The cardiac phase advances linearly from 0 to 2π during each R-R interval and is reset to 0 for the next cycle. The inspiratory phase spans from 0 to π and the expiratory phase spans from 0 to −π.

**Supplementary Figure 2**. Subject inclusion and exclusion.

**Supplementary Figure 3.** Distribution of brain lesions in the patient group (n=10). Lesions in most patients occurred in inferior frontal and anterior temporal areas.

**Supplementary Figure 4.** Examples of normal and poor quality plethysmography signals that could not be used for physiological correction. Arrows indicate the artifacts due to subject’s motion. Arrowheads indicate the loss of plethysmography signals.

**Supplementary Figure 5**. Brain connectivity derived following RETROICOR pipeline (upper panel) and bandpass filtering of 0.008-0.125Hz in BANDPASS pipeline (lower panel) in a representative patient and control subject. The connectivity links shown in the connectograms represent Pearson’s correlation coefficients >0.8 and exist after correcting multiple comparisons (p_fdr_<0.05).

**Supplementary Figure 6**. Spontaneous fluctuations at 1Hz or below in resting condition. (1) Lundberg (1960); (2) Lenfant (1967); (3) Chan et al. (2020); (4) Birn et al. (2006); (5) Wise et al. (2004); (6) Mayer (1876); (7) Obrig et al. (2000); (8) Chang et al. (2013).

## 8 AUTHOR CONTRIBUTIONS

YGB and BLE conceived and designed the study. WRS, DF, JEK, YGB and BLE performed the experiments. STC analyzed the data. STC, YGB and BLE interpreted results of the experiments and wrote the first draft of the manuscript. STC, WRS, DF, JEK, VN, YGB and BLE edited and revised the manuscript. All authors reviewed and approved final version of manuscript.

## 10 ACKNOWLEDGMENTS

We thank Karen Rich and the Massachusetts General Hospital MRI technologists for assistance with data acquisition. We also thank the nursing staff and respiratory therapists of the Massachusetts General Hospital Neurosciences ICU, Multidisciplinary ICU, and Surgical ICU. We are grateful to the patients and families in this study for their participation and support.

## 11 DATA AVAILABILITY STATEMENT

All relevant data without subject identifiers are within the manuscript. Individual imaging data with subject identifiers cannot be shared publicly because of institutional policies regarding data sharing and the protection of research subject privacy. The IRB protocols and the consent under which the subjects received imaging scans did not include language that permits the inclusion of their images in public data repositories. Data are available for researchers who meet the criteria from Massachusetts General Hospital IRB of MassGeneral Brigham HealthCare. Researchers seeking to utilize the de-identified imaging data from this manuscript should contact Drs. Brian Edlow and Yelena Bodien.

