## Supplementary figures and images for "Correcting Cardiorespiratory Noise in Resting-state Functional MRI Data Acquired in Critically Ill Patients"

### Supplementary Figure 1

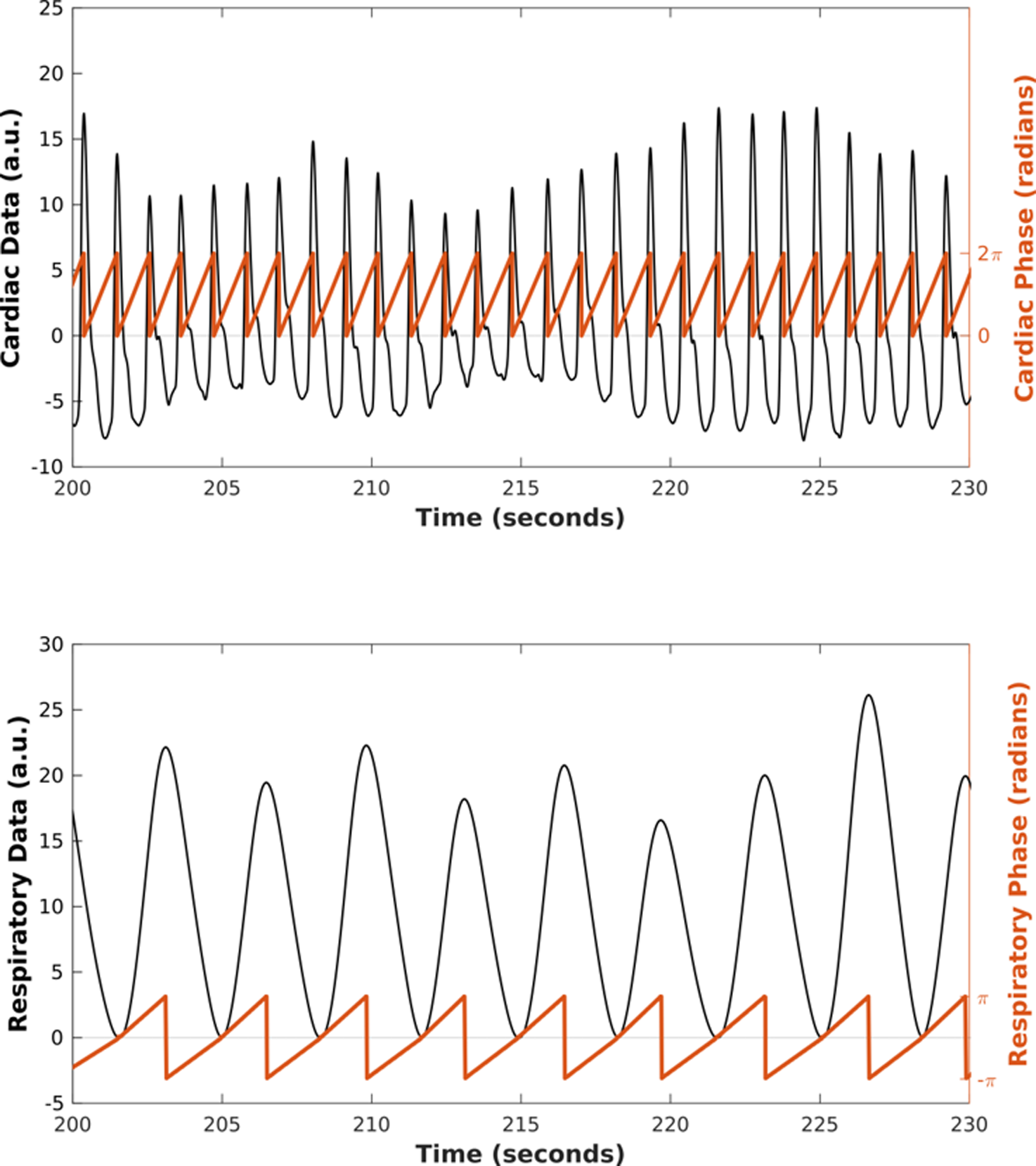

### Supplementary Figure 2

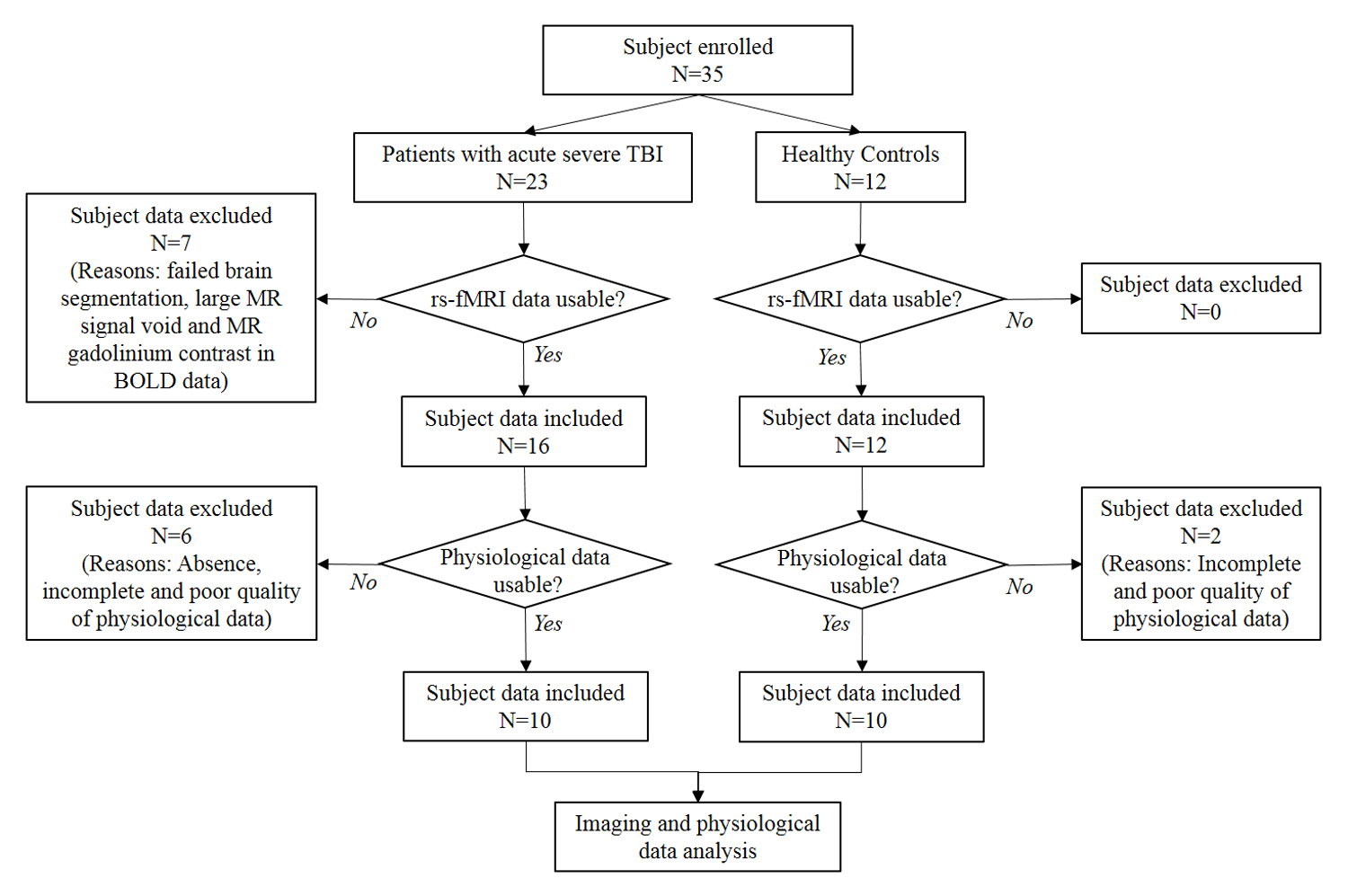

### Supplementary Figure 3

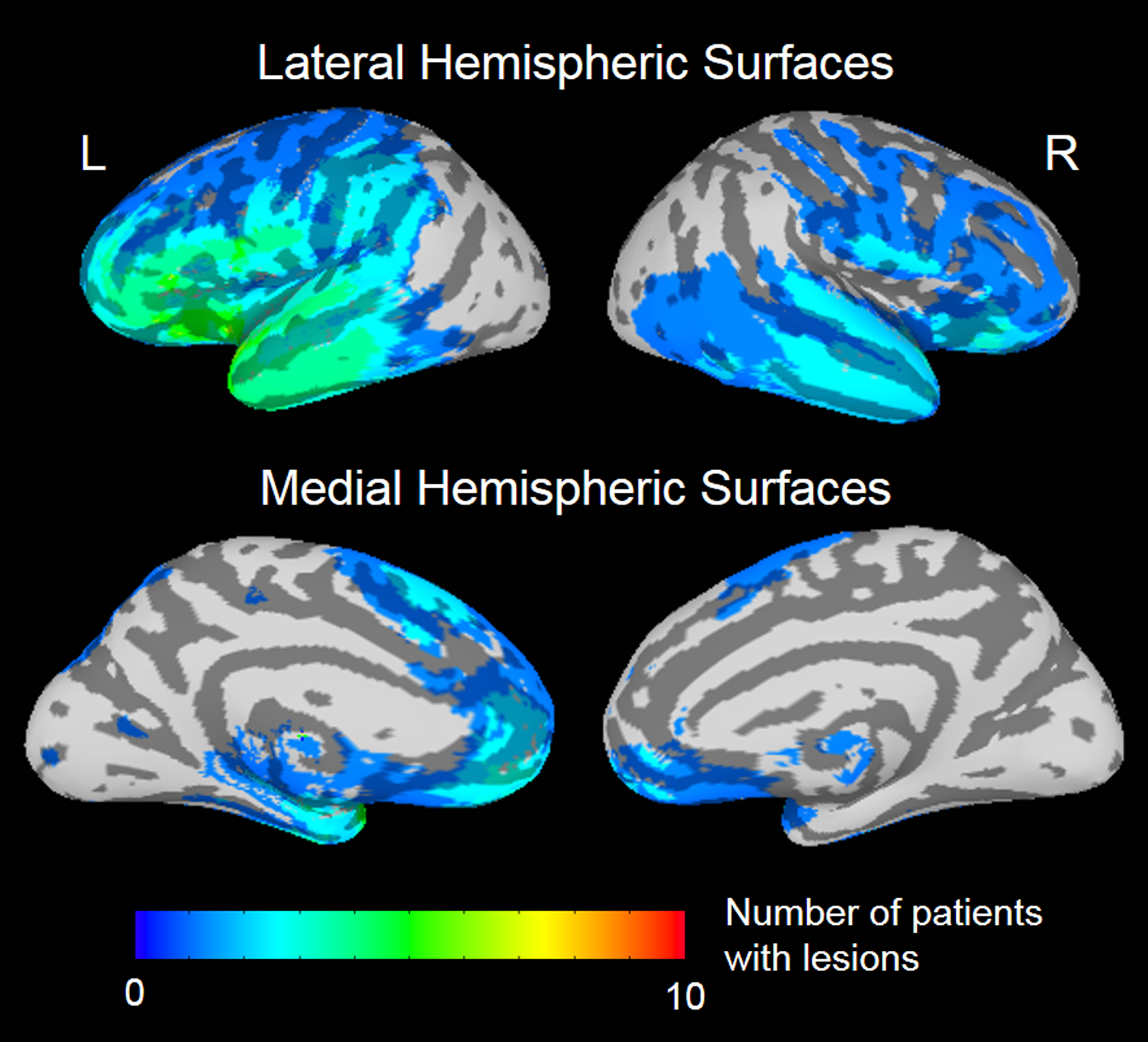

### Supplementary Figure 4

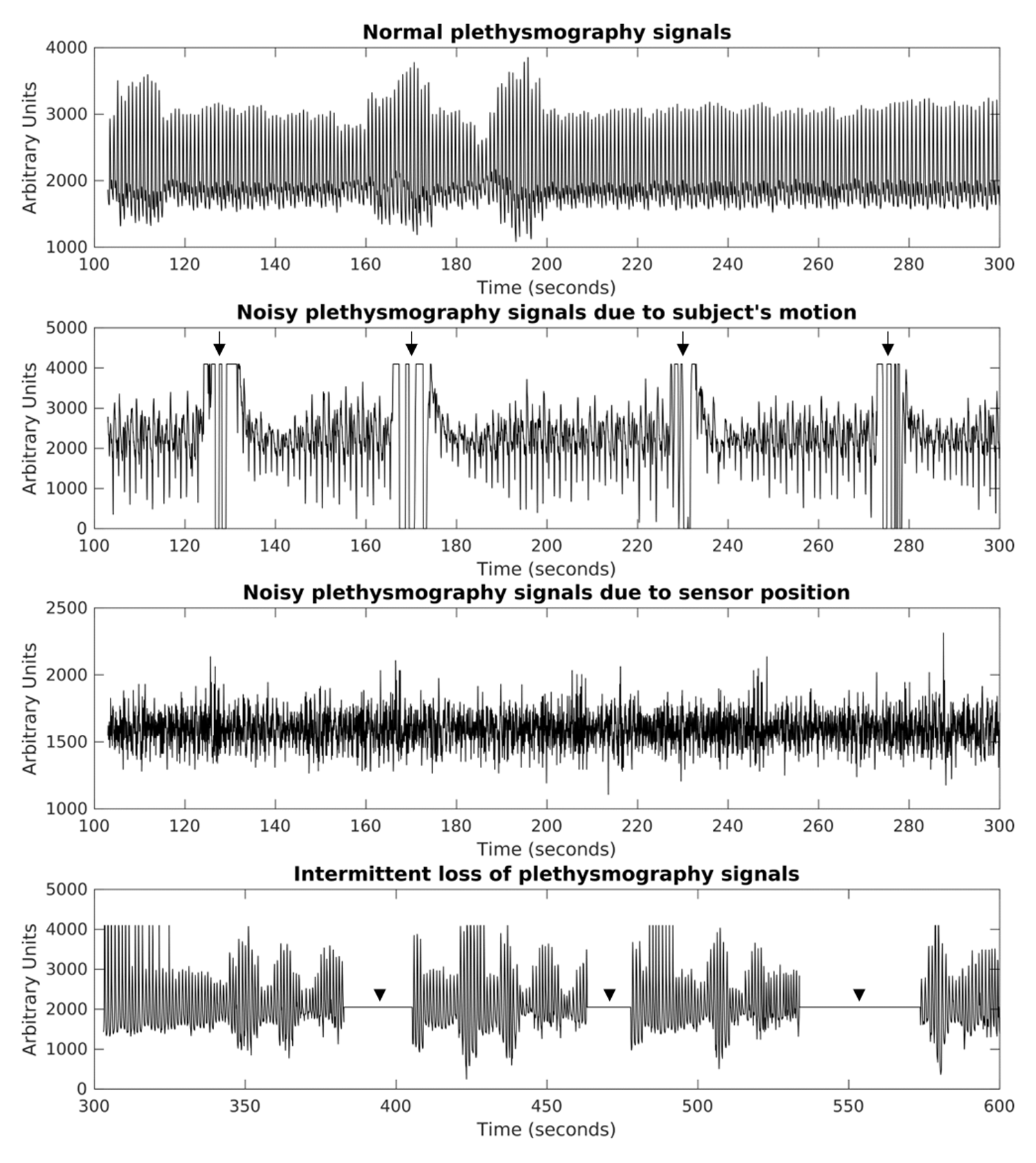

### Supplementary Figure 5

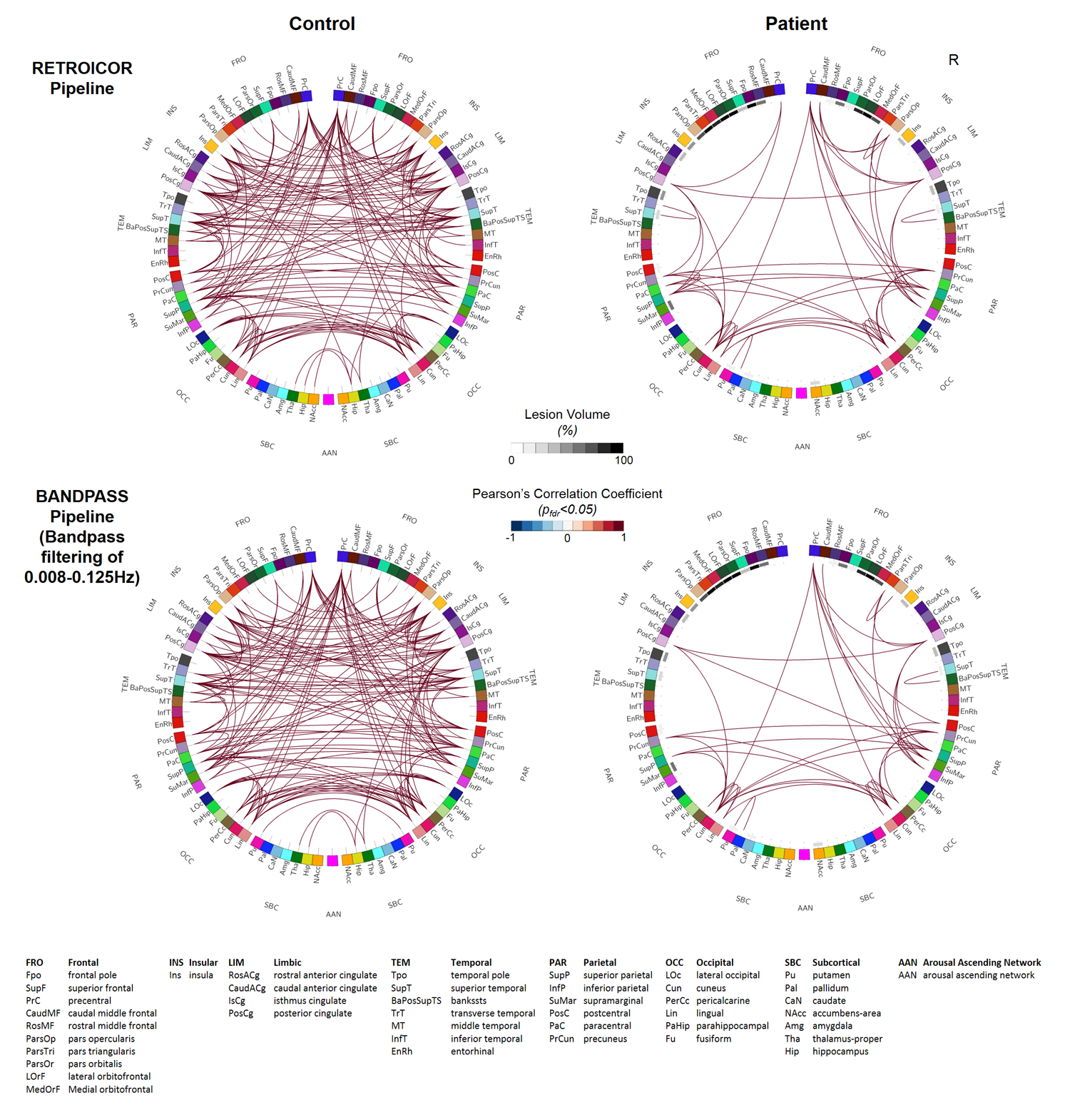

### Supplementary Figure 6

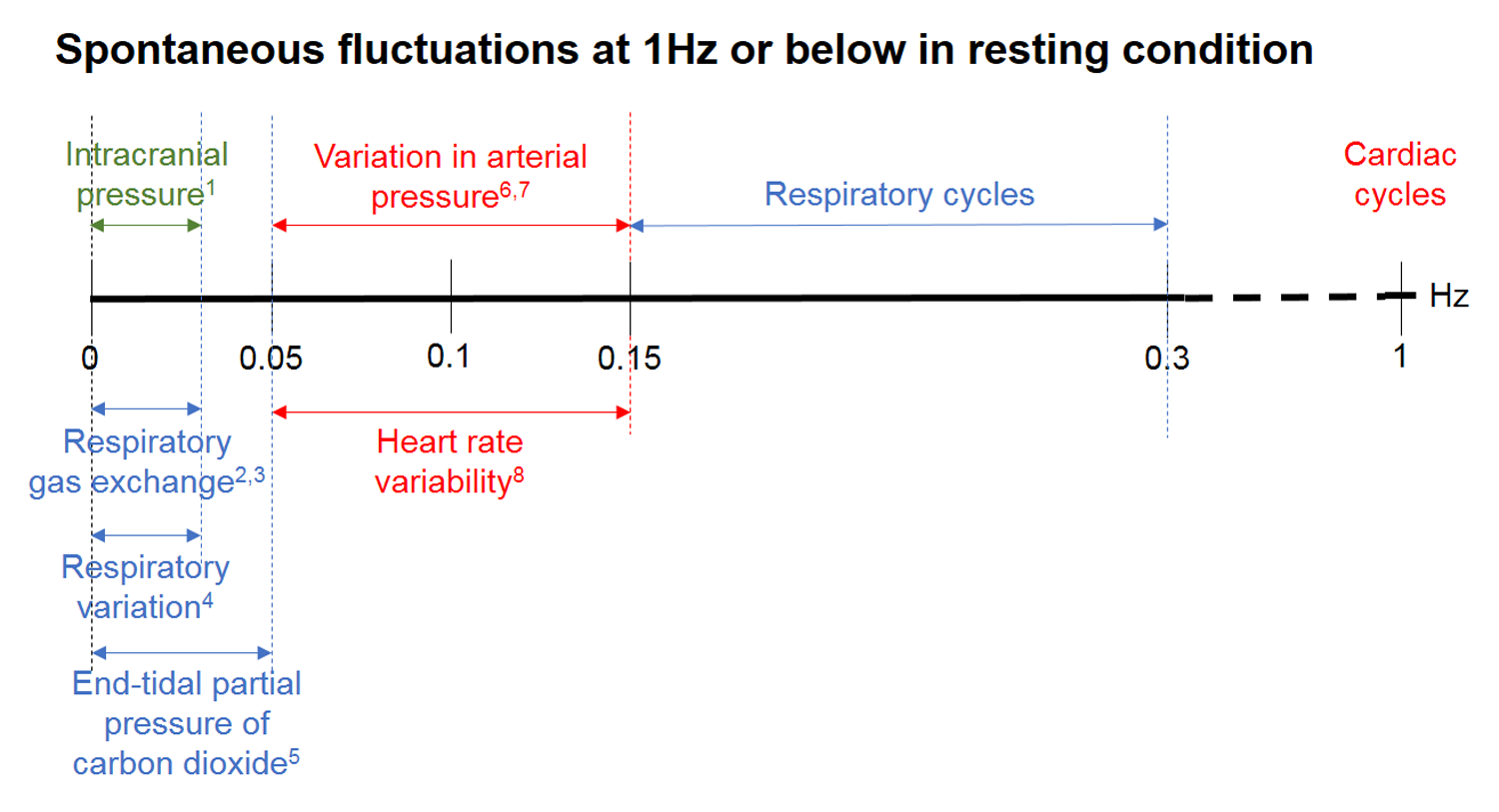
